# Mobile EEG reveals functionally dissociable dynamic processes supporting real-world ambulatory obstacle avoidance: Evidence for early proactive control

**DOI:** 10.1101/2020.09.16.298372

**Authors:** Magda Mustile, Dimitrios Kourtis, Simon Ladouce, Gemma Learmonth, David I. Donaldson, Magdalena Ietswaart

## Abstract

The ability to safely negotiate the world on foot takes years to develop in human infants, reflecting the huge cognitive demands associated with real-time dynamic planning and control of walking. Despite the importance of walking, surprisingly little is known about the neural and cognitive processes that support ambulatory motor control in humans. In particular, methodological limitations have, to date, largely prevented study of the neural processes involved in detecting and avoiding obstacles during walking. Here, using mobile EEG during real-world ambulatory obstacle avoidance, we captured the dynamic oscillatory response to changes in the environment. Time-frequency analysis of EEG data revealed clear neural markers of proactive and reactive forms of movement control (occurring before and after crossing an obstacle), visible as increases in frontal theta and centro-parietal beta power respectively. Critically, the temporal profile of changes in frontal theta allowed us to arbitrate between early selection and late correction mechanisms of proactive control: our data show that motor plans are updated as soon as an upcoming obstacle appears, rather than when the obstacle is reached, as previously thought. In addition, regardless of whether motor plans required updating, a clear beta rebound was present after obstacles were crossed, reflecting the resetting of the motor system. Overall, our use of mobile EEG during real-world walking provides novel insight into the cognitive and neural basis of dynamic motor control in humans, suggesting new routes to the monitoring and rehabilitation of motor disorders such as dyspraxia and Parkinson’s disease.

## Introduction

Moving safely through the environment while walking requires continual monitoring and adjustment of planned behaviour, including the ability to make fast online motor transformations in response to dynamic changes such as the appearance of unexpected obstacles. The skill of negotiating the constraints of the environment while walking is sufficiently complex that it develops slowly throughout infancy (Mowbay & Cowie, 2020) and is progressively lost in aging and motor impairments such as Parkinson’s disease (Holtzer et al., 2014; Peterson & Horak, 2016). The gradual reduction in cognitive resources and motor control that occurs with aging and disease means that it becomes increasingly difficult to respond effectively to obstacles that are encountered while walking. Indeed, falls associated with stumbling or tripping over objects represent a critical factor in the increased mortality rates that are seen for elderly and neurologic patients (Tinetti et al., 1988; Kovacs, 2005; Weerdesteyn et al., 2006). Given the complexity and fragility of the processes involved in walking, it is clearly important to identify the neural processes supporting cognitive control during walking and obstacle avoidance, generating new targets for clinical practice (Alexander & Hausdorff, 2008; Peterson et al., 2016).

Over the last decade, growing research interest in human ambulation has led to the extensive recording of EEG (the electroencephalogram) during active walking on treadmills (Petersen et al., 2012; Severens et al., 2012; Gwin et al., 2010, 2011; Gramann et al., 2011; Wagner et al., 2012; 2016; 2019; and Seeber et al., 2014, 2015). Recorded from electrodes placed on the scalp, EEG provides a non-invasive representation of oscillatory brain activity produced during task performance, allowing the identification of functionally dissociable cortical mechanisms that drive human behavior (Buzsáki & Draguhn, 2004). To date, EEG studies of walking have revealed the activation of a number of ‘prefrontal’ brain signals before approaching an obstacle, reflecting the recruitment of additional cognitive resources. For example, Haefeli et al. (2011) recorded EEG while participants walked on a treadmill, finding increased activity over frontal electrodes in response to an acoustic signal that warned of upcoming obstacles. Similar findings have been reported using mobile fNIRS. Maidan et al. (2018) reported a higher hemodynamic response over prefrontal sensors when participants had to prepare to step over unanticipated obstacles (compared to during normal walking), an effect that was independent of the size of the object. These findings have also been extended by a recent EEG investigation of walking on a treadmill (Nordin et al., 2019) that showed a power increase in low-frequency oscillations (i.e. ranging from 3-13 Hz) while participants were walking at different speeds and stepping over foam obstacles (appearing from behind a curtain placed at the front of the treadmill). These oscillatory brain changes were widespread across the scalp, consistent with activation of a distributed cortical network (i.e., supplementary motor, premotor and posterior parietal areas). Indeed, Nordin et al. (2019) argued that obstacle avoidance involves the early engagement of premotor and supplementary motor areas and a later activation of posterior parietal cortex.

Whilst studies of walking have focused largely on identifying neural markers, wider interest in the processes involved in goal directed behaviour have led to the development of theoretical models of cognitive control that provide a framework for understanding ambulatory control. Notably, studies of cognitive control by Braver and colleagues (Braver, 2012; see also Pezzullo & Ognibene, 2012) have characterised two broad stages of control processing. First, when a behaviour is planned, proactive control processes are employed to respond to potential sources of interference, allowing the original goal to be reached. Second, when an unexpected event has occurred, reactive control processes are employed to allow recovery from the interference, and return to the original goal. Markedly similar distinctions between proactive and reactive control mechanisms have also emerged from studies on human balance (Horak et al., 2006; Shumway-Cook & Woollacott, 2007; Bhatt et al., 2018). Proactive strategies are used to anticipate the loss of balance before it occurs (due to some source of interference), when the body has enough space and time to predict the upcoming interference and adjust motor plans. By contrast, reactive strategies involve compensatory adjustments that occur after unexpected events, to restore postural control and balance.

Although the theoretical distinction between proactive and reactive control strategies was not developed in relation to ambulatory control *per se*, the distinction is nonetheless clearly relevant for understanding the processes supporting obstacle avoidance during walking. Indeed, the neural signals observed in studies of treadmill walking can be readily interpreted within this theoretical framework. For example, Nordin et al. (2019) found widespread power modulation of low frequency oscillations, which was evident two steps before reaching unexpected obstacles, consistent with the operation of a proactive control process. According to this view, the putative proactive mechanism allows adjustments to be made to the planned walking activity. In addition, on the basis of the timing of the EEG signal, Nordin and colleagues proposed that the adjustment was made just before the obstacle was encountered, that would be suggestive of late correction.

To our knowledge there is no equivalent evidence of EEG markers of reactive control during obstacle avoidance. There is, however, wider evidence for reactive control mechanisms after movement. In particular, EEG studies have revealed post-movement increases of beta power (13-30 Hz), described as the beta rebound, as a marker of reactive control (Liebrand et al., 2017). Beta oscillatory activity over sensory motor regions is enhanced when the predictions of an incoming stimulus are violated (Arnal et al., 2011) and after forcibly interrupted movements (Alegre et al., 2008; Heinrichs-Graham et al., 2017), suggesting a mechanism that re-calibrates the motor system after a movement (Pfurtscheller et al., 1996; Engel & Fries, 2010; Kilavik et al., 2013). Thus, although reactive control mechanisms have not been demonstrated during obstacle avoidance, changes in beta power provides a likely candidate index of the operation of such mechanisms.

The recent emergence of mobile EEG (Ladouce et al., 2017) represents a particularly important development for researchers interested in walking, not least because it significantly extends the range of contexts in which brain activity can be studied (e.g., see Park & Donaldson, 2015; Park et al., 2018). Critically, using mobile EEG technology it is now possible to monitor the activity of the brain while participants navigate natural environments, taking investigations of walking off treadmills, and out of the laboratory (e.g., see Ladouce et al., 2019; Park & Donaldson, 2019, for examples). As a result, the neuro-cognitive processes supporting walking can now be studied in the real-world, offering an entirely new embodied perspective to the understanding of human behavior and motor impairments (which had been previously limited to non-ecological settings and fairly uninformative tasks; cf. McFadyen et al., 2017; Ladouce et al., 2017). Furthermore, the high temporal resolution of EEG (i.e., millisecond accuracy), combined with wireless portability, makes mobile EEG ideally suited to capturing the rapid cortical-responses that occur in response to dynamic stimuli (Kline et al., 2015).

As far as we are aware, currently there is no direct evidence for EEG markers of proactive and reactive control processes during real-world ambulatory obstacle avoidance. Thus, our primary aim in the current study is to ask whether the putative changes in EEG power described above can be identified during naturalistic walking. To address this issue we recorded EEG whilst participants walked across a room, to demonstrate whether it is possible to identify neural signals of proactive and reactive control during real-world obstacle avoidance. Rather than simply observe natural walking in isolation, however, we also examined EEG when obstacles were present. Critically, we manipulated the nature of the obstacle across trials, providing participants with more or less time and space to prepare for the obstacle. Obstacles were either absent, always present at the start of the journey, or appeared up ahead after a short or long delay. We highlight that our experimental design includes a condition in which no obstacle was presented - providing a baseline in which reactive and proactive control was not required. In addition, we manipulated the available time and space that participants had to adjust their gait when negotiating the environment, while allowing the walking task to remain as natural as possible. Based on the literature reviewed above, we predicted that proactive and retroactive control mechanisms should be identifiable in distinct temporal dynamics of theta and beta oscillations.

As well as demonstrating that neural markers of movement control can be identified during natural walking, we also tested two specific hypotheses. First, by varying the time and space that participants had to prepare for an obstacle we were able to arbitrate between early selection and late correction mechanisms of proactive control. As noted above, current evidence (cf. Nordin et al. 2019) suggests that proactive control mechanisms operate when an obstacle is tackled (so-called late correction). Here we test an alternative possibility, namely that proactive control processes operate as soon as information about an upcoming obstacle becomes available (so called early selection). Put simply, the high temporal resolution of EEG data would allow us to reveal the precise temporal dynamics of proactive control during walking. Second, by varying the opportunity to anticipate and prepare before adjusting to an obstacle, we aimed to test whether reactive control processes during walking are indexed by changes in beta power (the so-called beta rebound), reflecting the need for recovery after a change of a motor plan, in order to reset the previous state. Critically, based on previous studies (Arnal et al., 2011; Alegre et al., 2008; Heinrichs-Graham et al., 2017), we predicted that the beta rebound should only occur after participants cross an obstacle, and should be stronger when participants had less time to adjust their gait. As we show below, mobile EEG does indeed capture the dynamic engagement of proactive and reactive control processes during real-world ambulatory obstacle avoidance.

## Materials and methods

This study was approved by the local ethics committee and conformed to standards set by the Declaration of Helsinki. Thirty-two healthy participants (21 female and 11 male; age range = 19-65; mean age = 32.1 years, SD = 11.6 years) took part in the experiment. All participants were right handed (self-reported) and gave their written informed consent before the experiment.

The experimental design involved four conditions (as depicted in Figure 1) in which participants walked across an 18 m long room, passing through a series of infrared laser beams that recorded their location and controlled the presentation of obstacles (visible as a colour patch projected onto the floor that had to be stepped over). In the “no adjustment” condition no obstacle was presented and participants simply walked across the room. In the “preset adjustment” condition, obstacles were present at the start of each trial, placed at a fixed location 250 cm from the first laser beam. In the “immediate adjustment” condition, walking through the laser beam would trigger the presentation of an obstacle, displayed 160 cm in front of the participant. Finally, in the “delayed adjustment” condition, walking through the laser beam once again triggered the presentation of an obstacle, presented 310 cm in front of the participant. The participants were always instructed to walk straight across the room, to maintain a comfortable pace, and to step over any obstacle presented in front of them. Each crossing of the room corresponded to an individual trial, and on reaching the end of the room participants were asked to turn around and walk back across the room in the same way. The video projector and laser beams were arranged to allow data collection in both directions. Participants completed a total of 240 trials divided into 6 experimental blocks. Each block lasted for 5 min. All conditions were presented with equal probability. The overall experimental session lasted approximately 90 minutes, including preparation, recording and breaks between experimental blocks.

**Fig. 1.**
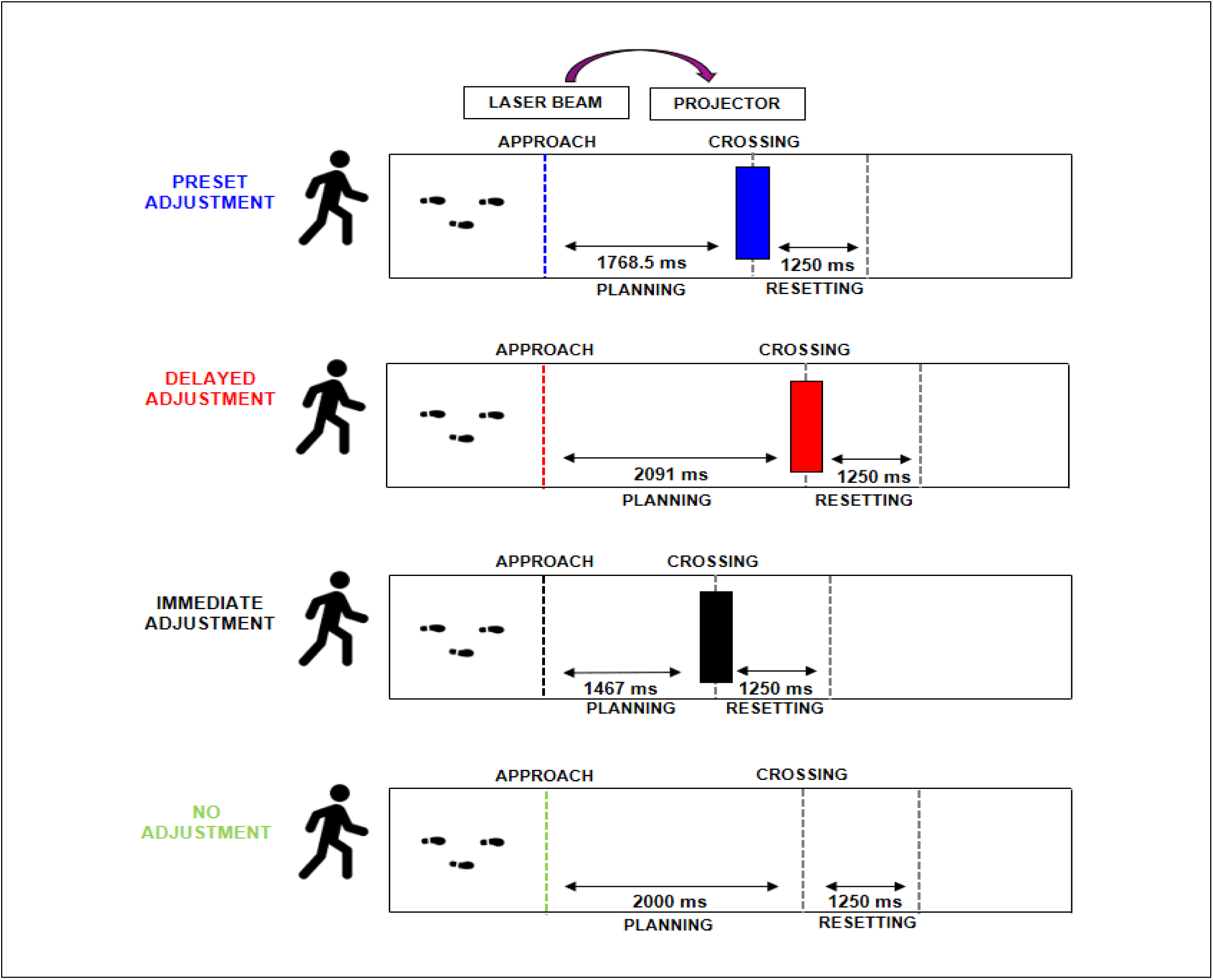
Representation of the experimental conditions, indicated with different colors (from top to bottom, respectively: blue, preset adjustment; red, delayed adjustment; black, immediate adjustment; green, no adjustment). For each condition, the median duration (in ms) of the planning phase (between participants) is reported inside each path between the approach and the crossing dotted lines.

The obstacle was presented as a white stripe (40×80cm) projected on a 10 m long carpet. The obstacle presentation was controlled with a system interfacing two fixed motion sensors placed at 230 cm from both ends of the carpet (directing infrared laser beams across the room, through which participants would pass). Stimulus presentation was controlled using E-prime 3.0 software (Psychology Software Tools, Pittsburgh, PA) and a projector. The motion sensors were designed to send an input signal to the stimulus presentation software running on a laptop, using the Auxiliary I/O port of a Chronos response device (Psychology Software Tools, Sharpsburg, PA). The laptop was connected to a projector placed at the side of the room. The presence and location of the obstacle presented varied on a trial-by-trial basis, depending on the experimental condition.

During each trial the experimenter manually marked two main events (as illustrated in Figure 1): the moment that the participant crossed the beam (‘Approach’) and the moment when the participant was over the obstacle (‘Crossing’). These two points provided temporal markers for use within the analysis of the EEG data that identified a planning phase (before the obstacle was encountered) and a resetting phase (after the obstacle was encountered).

Participants also wore foot sensor insoles (Pedar-x System, novel.de, Munich, Germany), a bluetooth pressure distribution measuring system for monitoring local loads between the foot and the shoe. The data of gait parameters were not recorded in all participants of this study and are not reported here.

### EEG acquisition and analysis

EEG data was recorded from 32 Ag/AgCl electrodes connected to a portable amplifier (ANT-neuro, Enschede, The Netherlands). Electrodes were positioned according to the International 10-20 system (FP1, FPz, FP2, F7, F3, Fz, F4, F8, FC5, FC1, FC2, FC6, M1, T7, C3, Cz, C4, T8, M2, CP5, CP1, CP2, CP6, P7, P3, Pz, P4, P8, POz, O1, Oz, O2) with AFz electrode as ground and CPz electrode as reference. The electrode impedances were reduced below 5 kΩ before the recording. EEG data were sampled at 500 Hz and bandpass filtered at 0.01-250 Hz.

EEG data analyses were performed using custom scripts written in MATLAB 2019a (The MathWorks) incorporating EEGLAB toolbox (Delorme and Makeig, 2004). Mastoid channels (M1 and M2) were removed from the initial 32 electrodes. EEG channels with prominent artifacts were automatically selected (kurtosis > 5 SDs) and interpolated. All channels were then re-referenced to the average. An extended infomax Independent Component Analysis (ICA, Makeig et al., 1996) was performed, using a 0.1 Hz to 40 Hz bandpass filter. After the ICA decomposition, a Finite Impulse Response (filtered from 0.1 Hz to 80 Hz, −6db cut-off, filter order 16500) was applied to ICs in order to nullify phase delay. Resulting non-artifactual ICs scalp maps were selected through SASICA (Semi-Automated Selection of Independent Components of the electroencephalogram for Artifact Correction, Chaumon et al., 2015). The selected ICs scalp maps, residual variance and mean power spectra were further visually inspected to identify non-brain sources. An average (mean ± SD) of 5.85 ± 1.97 of non-artifactual ICs across conditions were selected and retained for the analysis.

Data was segmented into epochs relative to the step over the obstacle (i.e., the ‘Crossing’ event, which was defined as time 0), producing a −3500 ms to 2000 ms time window. Since the latency of different trials were affected by a great deal of variability within and between participants, single trial spectrograms were time warped to the median latency (across participants) of the ‘Approach’ event using linear interpolation. In order to have the same number of trials, 40 trials were randomly selected for each condition. Epochs still contaminated by muscular artifacts or in which the latency of the ‘Approach’ event exceeded the limit of −2500 ms from the ‘Crossing’ were excluded. An average (mean ± SD) of 37 ± 2.07 epochs across conditions were included in the subsequent analysis, resulting in 7.5% of trials being excluded. Event related spectral perturbations (ERSPs) were obtained by computing the mean difference between single-trial log spectrograms for each channel, for each participant, relative to the mean baseline spectrum (from −3000 ms preceding to 1500 ms following the obstacle stepping).

## Statistical Analysis

Midline single channel spectrograms (Fc, Cz, POz; Figure 2) were visually inspected to identify prominent changes in the spectral power across conditions. Topographic maps of theta (Fig. 3) and beta (Fig. 5) activities were further assessed to determine the cortical origin of relevant spectral changes for each frequency band. Informed by our hypothesis and visual inspection of the topographic maps, we identified a frontal (mean activity across FC1, Fz and FC2 channels), a central (CP1, Cz and CP2) and a parietal (P3, POz and P4) location Finally, in order to examine the time course of spectral changes before and after the obstacle, the planning (from −1750 ms to −250 ms) and the resetting (from 250 ms to 1250 ms) periods were divided into a series of successive 500 ms time windows. Four different repeated measure ANOVAs with 3-within factors (experimental condition, time window and location) were performed to examine the pattern of activity across the planning and the resetting phases for each frequency band. Significance level was set at p < .05 and where sphericity assumption was violated the Greenhouse-Geisser method was used to correct the degrees of freedom. Post-hoc paired samples t-tests were adjusted for multiple comparisons using Bonferroni correction.

**Fig. 2.**
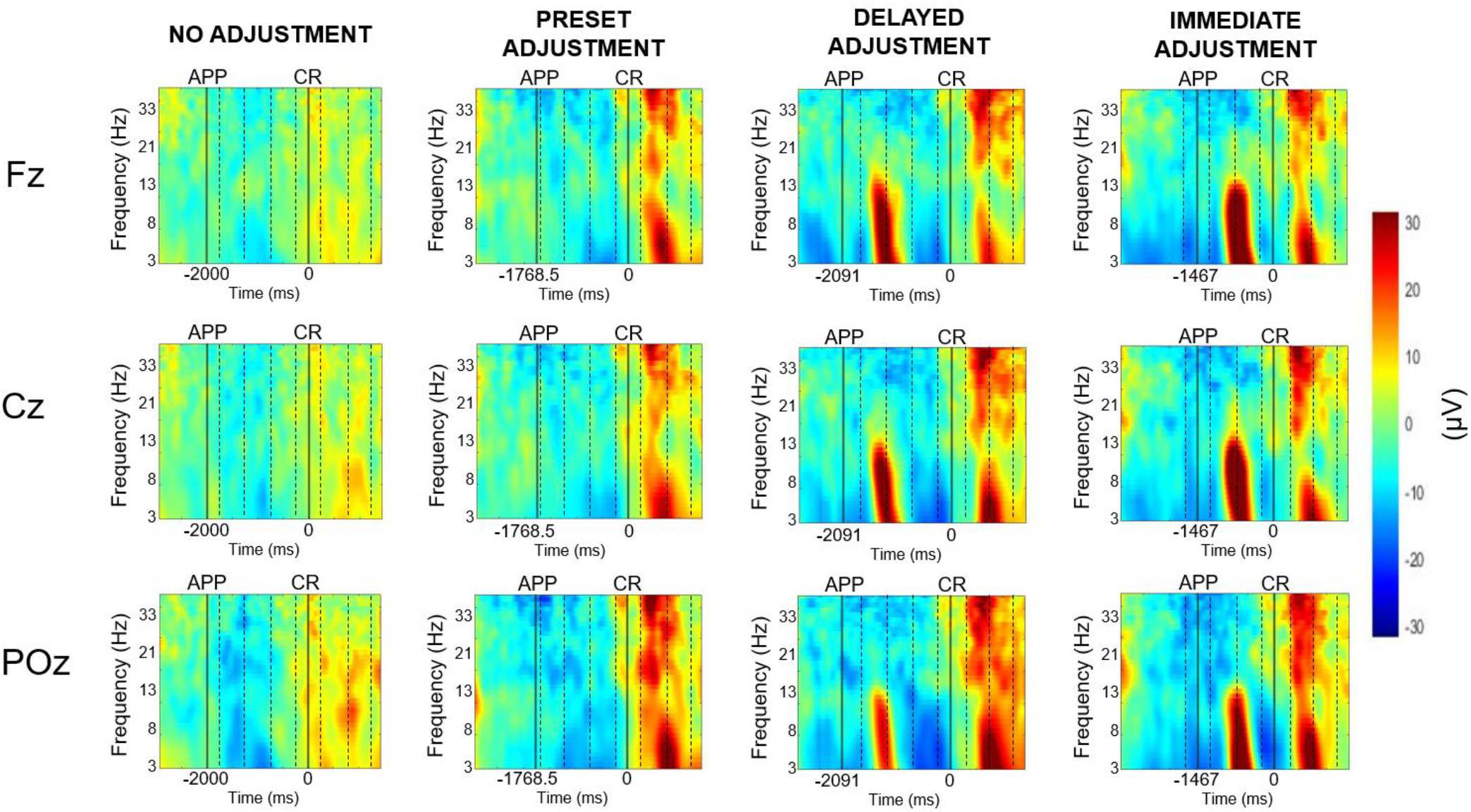
Time warped spectrograms at electrodes Fz, Cz, and POz for each experimental condition. Vertical solid black lines represent the ‘Approach’ (APP) and the ‘Crossing’ (CR, time 0) events, respectively. Vertical dotted lines represent time windows included in the analysis. On the x-axis (time in ms), the median latencies of the timing of the Approach point are reported for each condition. The lowest frequency shown is 3 Hz, the highest is 35 Hz. Colors indicate the relative change of power from the baseline. Blue colors represent decrease of power; red colors indicate increase of power.

**Fig 3.**
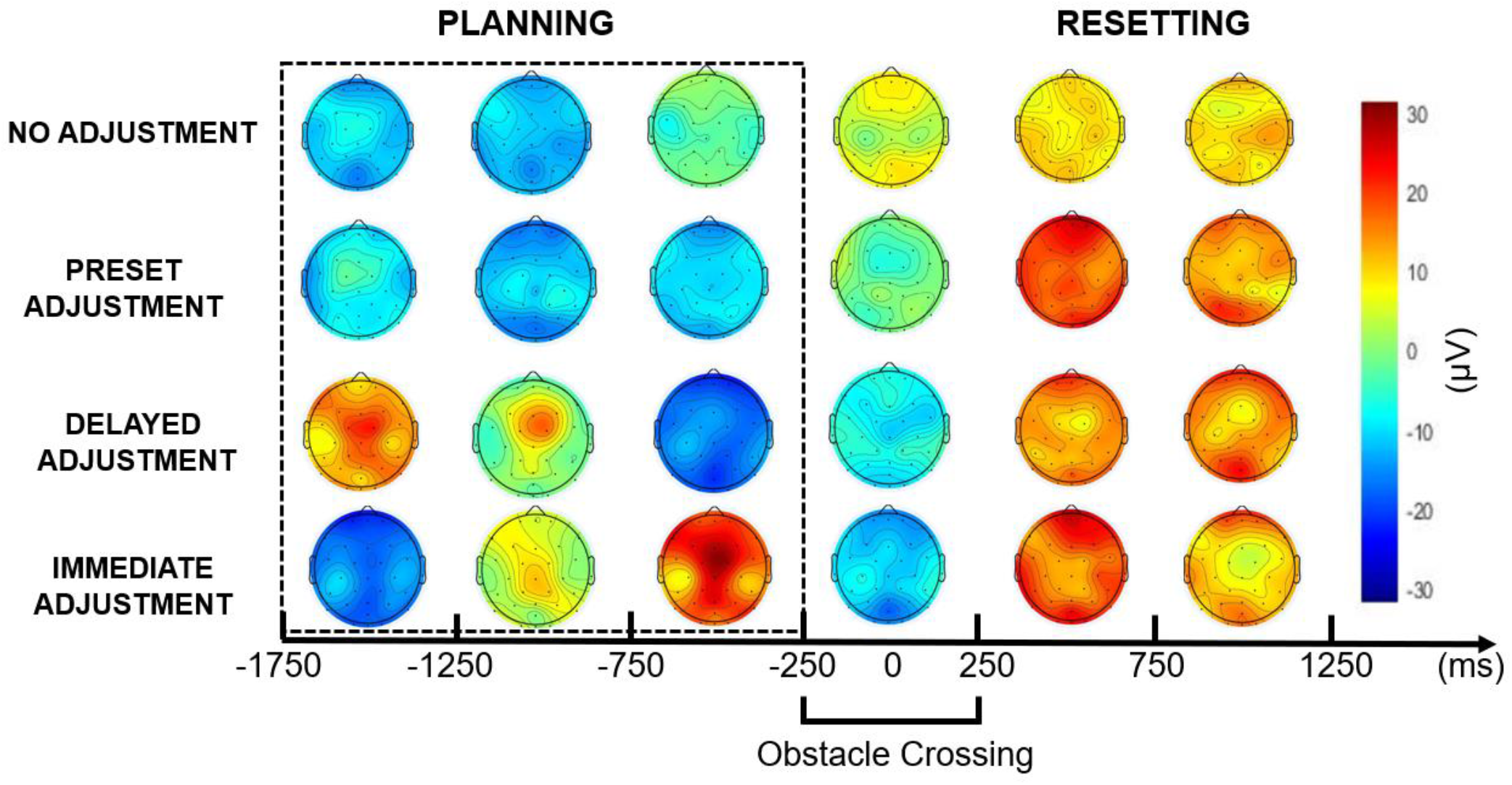
Topographic maps illustrating the temporal dynamics of theta activity across conditions and time windows. The dotted rectangle around the scalp maps before time 0, indicates the time windows included in the planning phase.

**Fig. 4.**
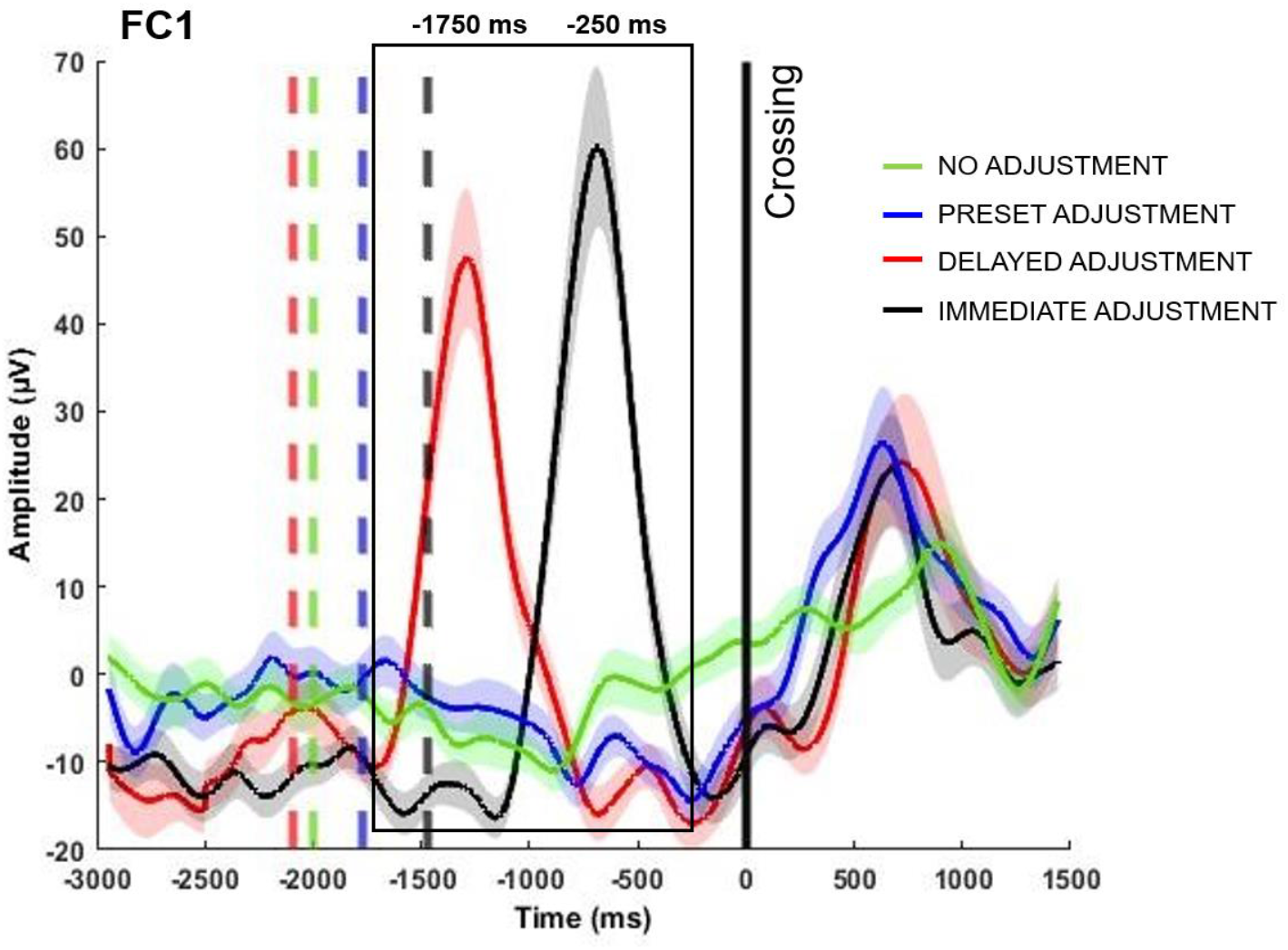
The time course of changes in theta across the experimental conditions (group mean, with standard errors indicated by shading) shown for a representative electrode (FC1). Dotted lines represent the median latency of the ‘Approach’ event, that matches the same color of the conditions indicated by the key. Solid vertical black line indicates the ‘Crossing’ event (time 0). The black rectangle indicates the time windows included in the analysis of the planning phase.

**Fig. 5.**
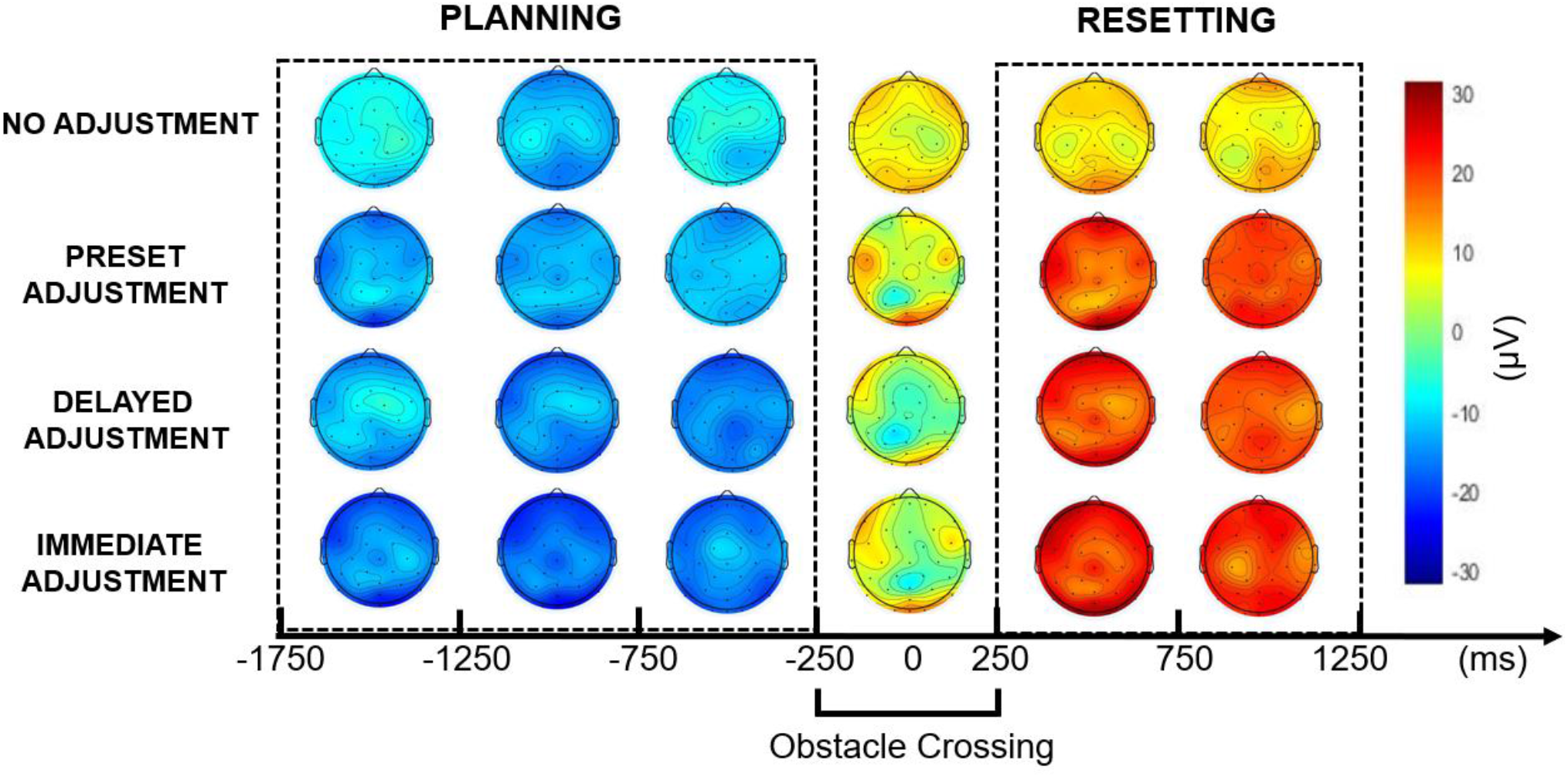
Topographic maps illustrating the temporal dynamics of beta activity across conditions and time windows. The dotted rectangles around the scalp maps before and after time 0, indicates the time windows included in the planning and in the resetting phase respectively.

## Results

Midline time warped spectrograms (Figure 2) revealed a transient change in the spectral power of theta (4-7 Hz) and beta (13-35 Hz)^1^ frequency bands, occurring after the ‘Approach’ and before ‘Crossing’ and differently distributed across conditions. Below the results for each frequency band will be presented separately for the planning and the resetting phases.

### Planning

Theta. The ANOVA indicated that changes in the theta spectral power were significantly different across experimental conditions [F(1, 31) = 14.645, p < .001, ῃ_p_^2^ = .321]. Post-hoc paired sample t-tests revealed that the theta increase was significantly stronger both in the immediate adjustment [immediate vs no adjustment: t(31) = 6.150, p < .001; immediate vs preset: t(31) = 5.374, p < .001; immediate vs delayed: t(31) = 2.142, p < .05] and in the delayed adjustment condition [delayed vs no adjustment: t(31) = −4.235, p < .001; delayed vs preset: t(31)=−2.811, p < .01], but similar in the preset adjustment and no adjustment conditions (p = .375). A main effect of location [F (1, 31) = 8.302, p < .001, ῃ_p_^2^ = .211] revealed that the increase in theta increase was more pronounced at frontal compared to parietal [t(31)= 3.733, p < .001] and central [t(31) = −2.154, p < .05] electrodes, and decreased strongly in parietal compared to central [t(31) = 2.138, p <.05].

A significant interaction between experimental condition and time window [F (1, 31) = 37.313, p < .001, ῃ_p_^2^ = .546; Fig. 2] confirmed that the timing of the increase in theta power was consistent with the appearance of the obstacle in the immediate and delayed adjustment conditions. As shown in Figure 3, a significant stronger theta increase occurred firstly in the delayed adjustment after the obstacle appeared [−1750 ms to −1250 ms; delayed vs no adjustment: t(31) = −6.007, p < .001; delayed vs preset: t(31) = −4.150, p < .001; delayed vs immediate: t(31) = −5.598, p < .001] and decreased more in the immediate compared to preset adjustment condition [t(31) = −3.248, p < .01]. In the following time window (−1250 ms to −750 ms) the theta increase became stronger in the immediate adjustment condition [immediate vs no adjustment: t(31) = 4.922, p < .001; immediate vs preset: t(31) = 4.432, p < .001] but was still present in the delayed adjustment condition [delayed vs no adjustment: t(31) = −6.052, p < .001; delayed vs preset: t(31) = −3.345, p < .01]. In the last time window the theta increase was stronger in the immediate adjustment condition [immediate vs no adjustment: t(31) = 5.902, p < .001; immediate vs preset: t(31) = 6.904, p < .001; immediate vs delayed: t(31) = 10.882, p < .001], but the decrease was stronger in the delayed adjustment condition [delayed vs no adjustment: t(31) = 7.586, p < .001; delayed vs preset: t(31) = 2.163, p < .05] and in the preset adjustment conditions [preset vs no adjustment: t(31) = 3.602, p < .001]. Post-hoc t-tests revealed no statistical differences between preset adjustment and no adjustment conditions during the first two time windows (p > .05) of the planning phase. No other main effect or interaction reached the statistical significance (p > .05).

Beta. Although the ANOVA did not show a main effect of condition, beta decrease of power occurred stronger in the immediate adjustment condition (mean = −9.69 ± 7.09 μV), followed by the delayed adjustment condition (mean = −9.08 ± 6.73 μV), the preset adjustment condition (mean = −8.34 ± 6.72 μV) and no adjustment condition (mean = −5.43 ± 7.28 μV). A main effect of brain locations [F (1, 31) = 4.183, p < .05, ῃ_p_^2^ = .119] revealed that a stronger decrease in beta frequency band occurred in central (mean = −8.43 ± 4.66 μV) and parietal (mean = −8.80 ± 5.03 μV), areas compared to frontal (mean = −7.19 ± 4.82 μV), although post-hoc paired sample t-tests showed only one statistically significant difference [parietal vs frontal: t(31) = 2.589, p < .05]. A significant interaction between time windows and conditions [F (2, 31) = 2.919, p < .05, ῃ_p_^2^ = .086; Figure 6] showed that beta decrease was significantly stronger in all obstacle conditions compared to no adjustment in the last time window [−750 to −250 ms; no adjustment vs immediate: t(31) = −2.876, p < .01; no adjustment vs delayed: t(31) = 4.997, p < .001; no adjustment vs preset: t(31) = 3.742, p < .001]. A significant interaction between brain locations and time windows [F (2, 31) = 4,595, p < .01, ῃ_p_^2^ = .129] revealed that firstly beta decrease was stronger in parietal areas [time −1750 to −1250 ms; parietal vs frontal: t(31) = 2.219, p < .05] but later (−750 to −250 ms) when the participants were approaching the obstacle, became stronger in central areas compared to frontal [t(31) = −3.395, p < .01] and parietal [t(31) =-3.475, p <.01] areas. No other main effect or interaction reached the statistical significance (p > .05).

**Fig. 6.**
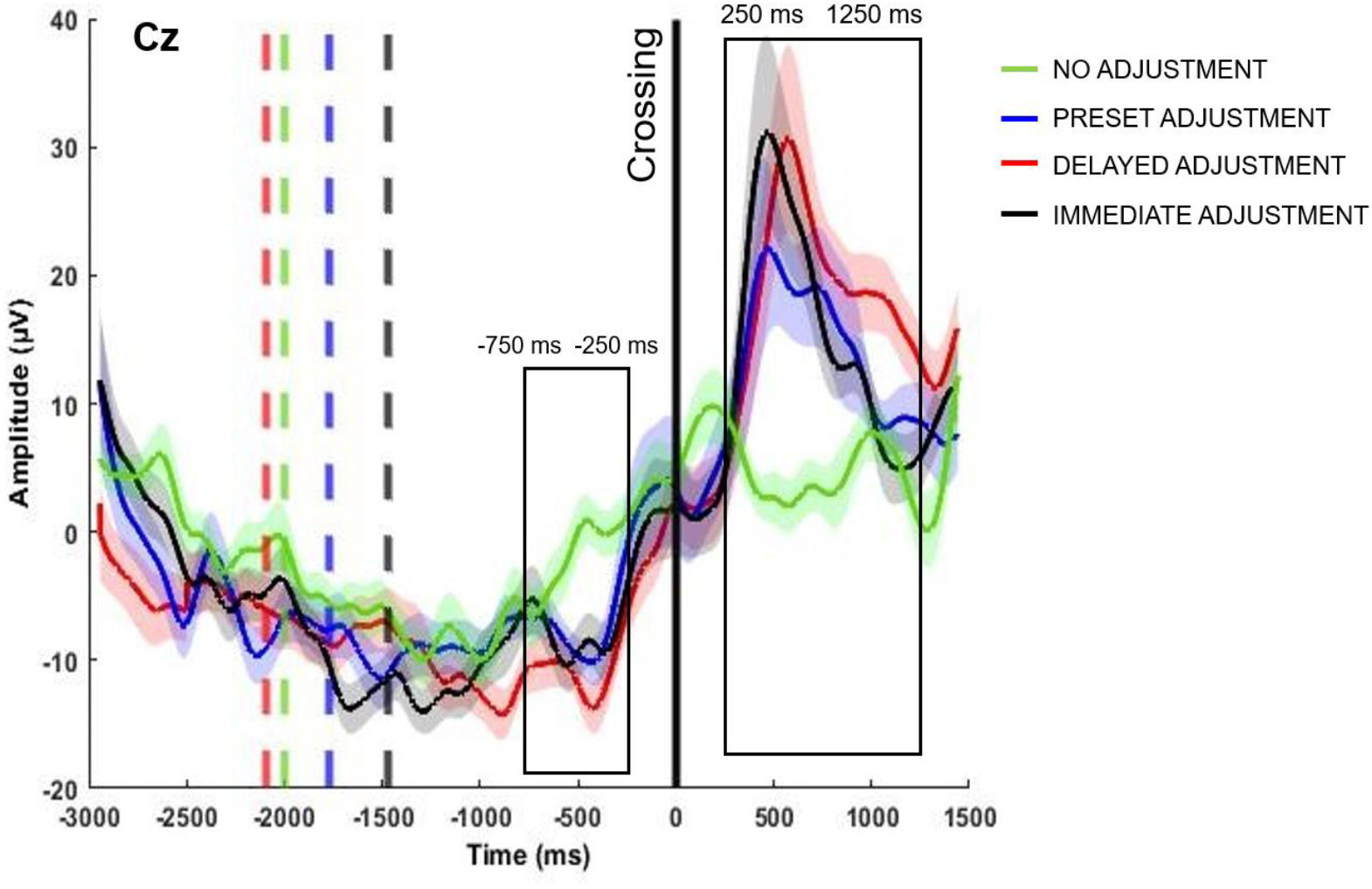
The time course of changes in beta across the experimental conditions (group mean, with standard errors indicated by shading) shown for a representative electrode (Cz). Solid vertical black line indicates the ‘Crossing’ event (time 0). The black rectangles indicate the time windows in which we found significant significant differences between conditions (−750 ms to −250 ms and 250 ms to 1250 ms respectively).

### Resetting phase

Beta. The ANOVA revealed a main effect of condition [F(1, 32) = 9.912, p < .001, ῃ_p_^2^ = .242] on beta modulation during the resetting phase. Beta increase of power was stronger in the all obstacle conditions compared to no adjustment [no adjustment vs immediate: t(31) = 4.525, p < .001; no adjustment vs delayed: t(31) = −5.113, p < .001; no adjustment vs preset: t(31) = −4.062, p < .001] condition. Additionally, beta increase of power was stronger in the delayed adjustment condition compared to the immediate adjustment condition [t(31) = −2.461, p < .05] but not compared to preset adjustment condition [immediate vs preprogrammed: p = .839; delayed vs preset: p = .258]. A main effect of brain location [F(1, 32) = 4.028, p < .05, ῃ_p_^2^ = .115] revealed that the beta increase was stronger in parietal compared to central [t(31) = −2.143, p <.05] and frontal [t(31) = −2.143, p < .01] areas. No other main effect or interaction reached the statistical significance (p > .05).

## Discussion

To our knowledge, this is the first mobile EEG investigation of real-world ambulatory obstacle avoidance. Our aim was to assess whether we can identify proactive and reactive forms of cognitive control during naturalistic movements, revealed through theta and beta spectral modulation, and to dissociate the cognitive processes involved.

The results showed greater transient theta spectral changes over frontal areas during the planning phase, consistent with the timing of the unexpected obstacles’ appearance on the path. The increase in theta power was larger when participants had less time and space available to change their gait before stepping over an obstacle (i.e. immediate adjustment condition). This pattern of modulation was substantially absent when participants walked without encountering any obstacle (i.e. no adjustment condition) or when they could see the obstacle in advance (i.e. preset adjustment condition). This novel investigation of real-world ambulatory obstacle avoidance, identifies increased theta as a marker of proactive cognitive control mechanisms when dealing with unexpected stimuli while walking.

The temporal dynamics of spectral power changes showed that the increase of theta power was linked to the appearance of the obstacle, suggesting the ‘early’ component of the proactive control is at play (Pezzullo & Ognibene, 2012) rather than a ‘late’, ‘just in time’ strategy (Braver, 2012). Nordin and colleagues (2019) investigated the brain dynamics during obstacle avoidance, and concluded that neural oscillatory activity in a similar frequency range (i.e. 3-13 Hz) increased two steps before the obstacle, pointing to a late correction mode of control. Instead, the present study shows that increases in frontal theta are not related to the time that an obstacle is tackled, but instead to the time that one becomes aware of an obstacle. Thus, the present results demonstrate that ambulatory obstacle avoidance relies on early selection models of proactive control.

According to the dual model theory, proactive control operates through mechanisms which maintain the relevant information actively in the brain until the behavior is accomplished (Braver, 2012). However, the continuous maintenance of goal-relevant information supporting complex behavior in the real world requires the recruitment of a large amount of cognitive resources. In situations where a planned action should not be immediately performed, proactive control ensures the flexible and cost-efficient updating of relevant information that ensures that the appropriate action will take place at the right time (Pezzullo & Ognibene, 2012). Indeed, recent evidence (Cooper et al., 2015; 2017, 2019) suggests that proactive control can be further divided in two stage-preparation processes: an early component, which ensure the preparation and the updating of relevant information to face a change, and a later component, that reflects motor readiness (Cooper et al., 2015, 2017, 2019). Our findings are a close fit with such a dual model account, as the brain dynamics in our study reveal a dissociation demonstrating that ambulatory avoidance of a partially unexpected obstacle relies on the earlier of the two stages of preparation processes. Increased frontal theta activity then serves an index of an ‘early’ proactive mechanism that prepares for an upcoming change, regardless of when the action is to be executed.

A stronger beta suppression during the planning phase was observed over sensorimotor areas only when the participants had to step over obstacles but not when there was no obstacle to avoid. As unobstructed walking involves a more basic negotiation of one’s environment, the greater beta band suppression over sensorimotor areas likely reflects a state of increased motor readiness, which is needed in order to negotiate the obstacle, without interrupting the walking cycle. Furthermore, we observed that beta suppression was initially stronger over parietal areas, but as the participants were closing on the obstacle the focus of beta suppression moved over central areas. Parietal beta suppression has been observed during visually guided step adjustments (Wagner et al., 2012) and motor programming of finger movements (Mars et al., 2007). The parietal cortex is associated with sensorimotor integration as well as spatial representation of ongoing movements (Buneo & Andersen, 2006; Andersen & Cui, 2009). Moreover, studies in cats (Drew et al., 2008; Drew & Marigold, 2015; Marigold & Drew, 2017) revealed neuronal populations involved in computing and approximating the time and space available to avoid contact with objects while walking. A central beta power decrease has been observed during active walking (Wieser et al., 2010; Presacco et al., 2011; Wagner et al., 2012; Seeber et al., 2014) and cycling (Jain et al., 2013; Storzer et al., 2016). It is well established that beta suppression over sensorimotor brain regions is an index of motor activation thought to reflect the planning and the execution of voluntary movements (Neuper et al., 2006; Pfurtscheller & Berghold, 1989; Pfurtscheller & Lopes da Silva, 1999). The temporal evolution of beta suppression in the present study points towards the operation of a sequential mechanism which initially recruits sensorimotor integration and spatial representation processes and at a later stage movement planning processes. Notably, the temporal evolution as well as the magnitude of beta suppression were similar when the gait adjustments were either preset or triggered by the presentation of the unexpected obstacle. This may be due to the relatively low difficulty of stepping over the obstacle in the present study and also suggests that updating of a motor plan, presumably reflected in theta increase, is not necessarily reflected in a greater activation of primary sensorimotor areas.

With regards to reactive control once the obstacle is tackled, and in addition to changes during the planning phase, we observed transient power changes in the beta and theta frequency bands also during the resetting phase. Consistent to our hypothesis, an increase in beta power, the so called post-movement beta rebound (Pfurtscheller et al., 2005; Pfurtscheller & Solis-Escalante, 2009; Jurkiewicz et al., 2009) was present only when gait adjustments were required in order to step over the obstacle, but absent when there was no obstacle to avoid. The beta rebound is typically observed generally over sensorimotor areas after motor execution or motor imagery (Pfurtscheller et al., 2005; Pfurtscheller & Solis-Escalante, 2009) and it is believed to reflect an active recalibration process that takes place after a change in the state of the motor system (Pfurtscheller et al., 1996; Engel & Fries, 2010; Kilavik et al., 2013). Notably, the beta rebound over prefrontal and sensorimotor areas is considered a possible index of reactive control (Cooper et al., 2019; Liebrand et al., 2017). Accordingly, the presence of the beta rebound in our study only after a gait adjustment was required in order to step over an obstacle, provides a marker of the reactive process that needs to take place in order to restore the motor system to its previous state.

We also predicted that the beta rebound should be stronger when participants had less time to adjust their gait. However, although the beta rebound was clearly present after tackling the obstacle, this index of recovery was not more pronounced when obstacles appeared while walking compared to when the obstacle was present at the start of the journey. Therefore, the beta rebound as an index of recovery is not modulated by the cognitive demands placed when adapting gait with limited time and space to do so. Interestingly, the beta rebound was prolonged when the participants had more time to adjust their gait before stepping over the obstacle. A recent study has suggested that the duration of the beta rebound is increased after temporally protracted movements (Fry et al., 2016). Accordingly, the prolonged beta rebound could be related to the longer engagement of the motor system when a partially unexpected obstacle appeared at a greater distance from the participants.

## Practical implications of the current study

An objective of the present study was to demonstrate the relevance and utility of using mobile EEG in real-world investigations, that allows the detection of neural correlates of human natural behavior that cannot be captured in the traditional laboratory settings, as we proposed already (Ladouce et al., 2017; 2019). Our data revealed the neuro-cognitive indices of proactive and reactive control during natural human ambulation, indicating that control of walking requires a sophisticated system emerging when negotiating complex real-world dynamics. Despite extensive development of new hardware solutions (i.e. dry electrodes, Lopez-Gordo et al., 2014, or dual-layer EEG caps, Nordin et al., 2019) and tools for signal processing (i.e. independent component analysis, Makeig et al., 1996) mobile technologies have not typically been utilised to provide exhaustive explanations of the human cognition of moving in natural contexts. The technical challenges of reducing motion artifacts during natural movements and the physical constraints of multiple portable devices, are still demanding much attention as these continue to present significant barriers for neuroscientific research. However the validity of our data is encouraging, and demonstrates a readiness for example for enhancing the use of the Mobile Brain/Body Imaging (MoBi) approach (Makeig et al., 2009) by combining the recording of neural activity through the mobile EEG with body signals (i.e. muscle activity, kinematics and eye movements) through portable devices. Using this approach, EEG brain activity is time-stamped with behavioral indexes, like heel strikes while walking. Although previous MoBi investigations provided significant insights on the coupling between gait rhythms and intracortical patterns (Severens et al., 2012; Gwin et al., 2010; Gramann et al., 2011; Wagner et al., 2012, 2016; Seeber et al., 2014, 2015; Bruijn et al., 2015), our study aimed to understand the nature of cognitive processes that guide behavior in real-world environments, rather than explaining the relation between gait parameters and cortical activity. Importantly, our goal was to highlight that the mobile approach of studying human behavior in real world environments may have important implications for the understanding of neurodegenerative disorders, such as Parkinson’s disease, and developmental disorders such as dyspraxia. Negotiating obstacles in the real-world requires us to allocate attention, detect relevant constraints and flexibly adapt motor behaviors, which is challenging for elderly or Parkinson’s disease patients who usually experience gait impairments that increase the risk of falls and mortality (Tinetti et al., 1988; Kovacs, 2005; Weerdesteyn et al., 2006). Studies that aimed to identify neural markers of Parkinson’s disease and gait dynamics are still limited, being restricted to simple tasks (i.e. finger tapping, Stegemöller et al.. 2016, 2017) or to kinematics recording (Vitório et al., 2010; Galna et al., 2010).

## Conclusion

Our study demonstrates that mobile EEG can be reliably utilised to capture the dynamic oscillatory responses associated with the neuro-cognitive indices of negotiating real-world environments. We demonstrated that naturalistic obstacle avoidance is mediated by proactive and reactive cognitive processes, reflected in the dynamics of theta and beta oscillations over frontal and sensorimotor brain areas. Temporal brain dynamics of frontal theta revealed that proactive control during unexpected obstacle avoidance is mainly supported by an early selection mechanism. Furthermore, we showed that motor readiness is mediated by beta power suppression over sensorimotor areas, that was present when preset or externally triggered gait adjustments were needed in order to step over an obstacle. With regard to reactive control, we identify a robust beta rebound after crossing an obstacle, demonstrating that real-world negotiation of the environment requires finely-tuned resetting of the motor system. What this shows is that by using mobile EEG, neuro-cognitive processes supporting walking can now be studied in the real-world, offering an entirely new embodied perspective to the understanding of human behavior and motor impairments and benefit the innovation of neuro-rehabilitation approaches.

## Acknowledgements

We would like to thank technicians Chris Grigson, Stephen Stewart and Catriona Bruce for their help with the experimental setup and recordings.

## Author Contributions

Author contributions: M.M., S.L., D.K., G.L., D.I.D. and M.I. conception and design of the study; M.M. performed the experiment; M.M. and S.L. analyzed data; M.M., S.L., D.K., D.I.D. and M.I. interpreted results and all authors contributed to the manuscript.

## Data Accessibility

On acceptance of the manuscript we will share both analysis pipeline scripts and the data and related information on the Open Science Framework.

## Abbreviations

EEG: electroencephalography
fNIRS: functional near-infrared spectroscopy
ICA: independent component analysis
I/O: input/output
SASICA: Semi-Automated Selection of Independent Components of the electroencephalogram for Artifact Correction.

## Footnotes

1 Analysis of alpha (8-12 Hz) oscillations is not included in this manuscript in order to focus on the predictions examined through analysis of the theta and beta bands. However for interest, alpha oscillations mirror the patterns of theta oscillations that we report here.

